# Sparse balance: excitatory-inhibitory networks with small bias currents and broadly distributed synaptic weights

**DOI:** 10.1101/2021.02.26.433027

**Authors:** Ramin Khajeh, Francesco Fumarola, LF Abbott

## Abstract

Cortical circuits generate excitatory currents that must be cancelled by strong inhibition to assure stability. The resulting excitatory-inhibitory (E-I) balance can generate spontaneous irregular activity but, in standard balanced E-I models, this requires that an extremely strong feedforward bias current be included along with the recurrent excitation and inhibition. The absence of experimental evidence for such large bias currents inspired us to examine an alternative regime that exhibits asynchronous activity without requiring unrealistically large feedforward input. In these networks, irregular spontaneous activity is supported by a continually changing sparse set of neurons. To support this activity, synaptic strengths must be drawn from high-variance distributions. Unlike standard balanced networks, these sparse balance networks exhibit robust nonlinear responses to uniform inputs and non-Gaussian statistics. In addition to simulations, we present a mean-field analysis to illustrate the properties of these networks.

## Introduction

A typical cortical pyramidal cell receives thousands of excitatory inputs [1] that, without the influence of inhibition, would drive extremely high firing rates. It has been suggested that the inhibition that moderates these rates sets up a balanced condition that causes neurons to operate in a regime where fluctuations, not the mean, of their inputs drive spiking, resulting in irregular sequences of action potentials [2–5]. A number of theoretical models have been developed to address E-I balance and the irregular firing of cortical neurons (see [6] for a review). In one class of balanced models [7, 8], the input to each neuron has three strong components – recurrent excitation, recurrent inhibition and feedforward excitation. These balance automatically as part of the network dynamics, leaving residual fluctuations that drive neuronal firing at reasonable rates. Although the presence of strong excitation and inhibition is compatible with the data, there is no evidence for the strong feedforward inputs required in these models [9], and some evidence against them [10–13]. For this reason, we examine the consequences of removing strong feedforward input in balanced models.

In balanced models, synaptic strengths are drawn independently from two probability distributions, one for excitation and another for inhibition. For standard recurrent models to generate spontaneous irregular (chaotic) activity, the synaptic weight distributions must have a variance of order 1*/K*, where *K* is the in-degree, i.e., the number of synapses per neuron [8, 14–16]. The excitatory (inhibitory) distributions are only non-zero for non-negative (non-positive) values, and typically their mean is of the same order as the square-root of their variance, with both being of order 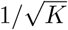. Summing this mean over the *K* synapses to each neuron gives a total input, which for reasons of stability is inhibitory, of order 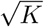. This large mean input must be cancelled and, in conventional models, this is done by adding constant feedforward excitation that is also of order 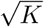. This is the large feedforward input that we aim to avoid.

A first question to ask is what happens if the order 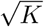 input is simply left out of the standard models, and replaced by an input of order 1. This results in a constraint on the firing rates; specifically, the average firing rate in the network must be of order 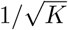. This implies that, although there can be irregular spontaneous activity without strong feedforward input, it involves neurons firing at very low rates. One way around this problem is to note that a small average firing rate is not incompatible with having individual neurons with significant firing rates if the activity is sparse. In other words, the average rate can be of order 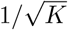 if, as in the standard models, activity is dense and individual rates are of order 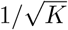, or if individual rates are of order 1 and the fraction of active neurons is of order 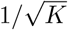. Here, we explore this latter possibility.

We mentioned above that standard balanced models require synaptic distributions with variance of order 1*/K* to generate irregular spontaneous activity. More precisely, the requirement is that, for each neuron, the sum of the variances of the strengths of its active inputs must be of order 1. In the standard model, this is satisfied because the product of *K*, the order of magnitude of the number of active inputs, and 1*/K*, the variance per synapse, is 1. In the sparse models proposed in the previous paragraph, the number of active inputs is only of order 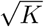, so the total variance computed in this way would be 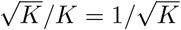, which is not sufficient to generate irregular activity. To solve this problem, we consider distributions of synaptic strength with means of order 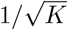, as in the standard model, but with much larger variances of order 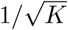. In this case, the total variance is of order 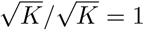, and irregular activity is restored.

Another feature of standard balanced models that seems at odds with the data is that they have attenuated linear responses to input that is uniform across neurons [9, 17]. This linearity is not present in the networks we study. In summary, the combination of small feedforward inputs and broadly distributed synaptic strengths gives rise to a novel E-I regime that exhibits asynchronous irregular sparse firing. In the following, we illustrate prominent features of this regime, such as nonlinear response to feedforward input and non-Gaussian current distributions, and we highlight the mechanisms that maintain sparsity and distributed firing across network neurons.

## Results

### The model

A common simplification for analyzing E-I networks is to consider a single population of inhibitory units driven by excitatory input from an external source [15, 16]. After analyzing such purely inhibitory networks, we will show that our results apply to networks with both excitatory and inhibitory units. We consider standard ‘rate’ models. The inhibitory networks we study have currents *x_i_* for *i* = 1, 2, …, *N* and firing rates *ϕ*(*x_i_*) that obey

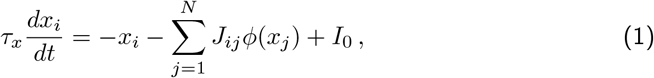

where *ϕ* is a nonlinear function and *J_ij_* ≥ 0. We call the variable *x* a current because it represents the total current generated by the recurrent synaptic and feedforward inputs in the second and third terms on the right side of the above equation, and because it determines the firing rate through the ‘F-I’ function *ϕ*(*x*). We also use the terms ‘response’ or ‘firing rate’ for *ϕ*(*x*) and ‘activity’ for non-zero rates. In our plots, we measure time in units of *τ_x_*, making it a dimensionless variable. Connectivity can be all to all (*K* = *N*) or we can restrict the connectivity so that only *K* < *N* of the elements in each row of *J* are non-zero. *I*_0_ is a positive bias input that is identical for all units; it is the feedforward input discussed in the Introduction. Standard balanced models assume the unrealistically large scaling 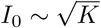; we consider, instead, models with *I*_0_ of order 1.

The non-zero elements of *J* are drawn independently from a distribution with mean 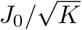, with *J*_0_ an order 1 parameter. We express the variance of this distribution as *g*^2^*/K^ν^*, where *g* is another parameter of order 1, and *ν* allows us to vary the scaling with *K*. The standard scaling is *ν* = 1, which we call low variance. As we will show, the novel sparse balance regime we explore comes about from setting *ν* = 1/2, which we call high variance.

It is awkward to use clipped Gaussians for sign-constrained synapses, especially in the large-variance case we consider. We use, instead, distributions with positive support, such as lognormal and gamma, focusing particularly on gamma-distributed synapses for reasons given below. This specific choice is not essential; network behavior remains qualitatively the same across a range of weight distributions, including a binary distribution (Fig S1).

We begin (Fig 1) by setting the response nonlinearity *ϕ* to a rectified hyperbolic tangent,

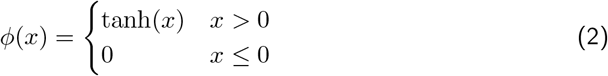

but later we also consider

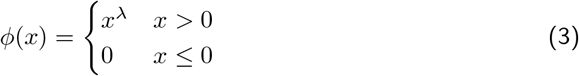

focusing, in particular, on the case *λ* = 0 (Heaviside function) for the analysis, but we also consider *λ* = 1 (rectified linear), and *λ* = 2 (rectified quadratic).

**Fig 1.**
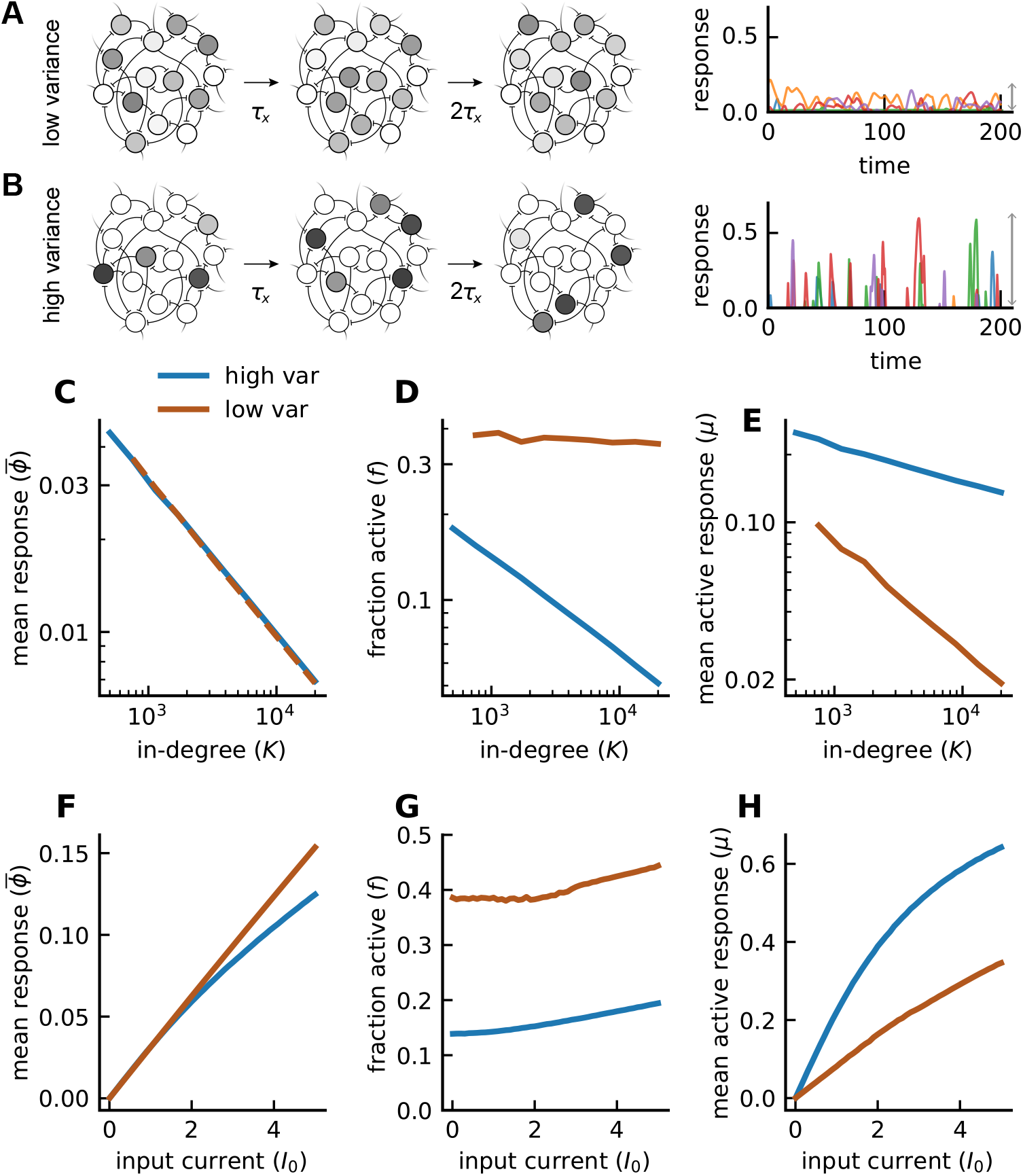
Comparison of low- and high-variance networks. **A)** Cartoon of network dynamics in time. Light (dark) gray corresponds to low (high) firing rates. With low synaptic variance, fluctuations in firing rates are small, and a relatively fixed and dense subset of units contribute to firing. Right: firing rate traces of five example units (each with a distinct color). Gray arrow indicates the extent of fluctuations in the network. **B)** Same as **A** except for the high-variance model. The network exhibits a small and shifting ensemble of cells that respond robustly at any given time. The magnitude of fluctuations is increased substantially (right). **C)** Mean response in both networks follows a 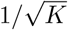 scaling (fits to the data yield 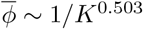 for high variance and 1*/K*^0.513^ for low variance; *J*_0_ is adjusted so that 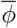 values in the two networks overlaps). **D)** Fraction of active units (inverse sparsity). High-variance model exhibits a rapid sparsening in *K* while, in the low-variance network, this fraction remains roughly constant. **E)** Mean response of the active subset. The trend in **D** is flipped: the low-variance network demonstrates a rapidly vanishing *μ*, which is not the case in the high-variance model. Input current *I*_0_ is set to one. **F)** Network’s response to the external input current *I*_0_ with *K* = 1000. **G)** Despite similar 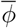 values, the high-variance network is more sparsely active by more than twofold. **H)** Active neurons respond more robustly in the high-variance network than in its low-variance counterpart. (Model parameters: *J*_0_ = 2 for high variance and 1.05 for low variance, *g* = 2, *J_ij_* ~ gamma, *ϕ* = [tanh]_+_).

Throughout, [·] denotes averages over units, 〈·〉 denotes averages over time, and an overline represents averages across both units and time. For fixed *K*, the results we present are independent of network size *N*, provided that the networks are large enough. For this reason and because we are interested in large *K*, we restrict our studies to the case *K* = *N*, but the results reported extend to partially-connected networks as well (*K* < *N* ; Fig S2).

### Simulation results

With the usual 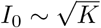 bias reduced to an input of order 1, the network behaves very differently in the low- (*ν* = 1) and high- (*ν* = 1/2) variance cases (Fig 1). For low variance, many units are active, but their responses are small (Fig 1A). In contrast, for high variance, activity in the network is sparse but individual units exhibit robust responses (Fig 1B). Scaling of the firing rate as a function of connectivity *K* can be quantified by computing

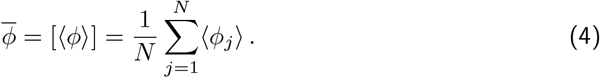

We can break down this average by writing it as the product of *f*, the fraction of units that are active (*ϕ* > 0), and *μ*, the average firing rate of the active units, 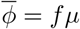. In both the low- and high-variance cases, the average firing rate 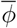 scales as 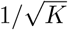 (Fig 1C) but, for low variance, *f* is fairly independent of *K* (Fig 1D) and *μ* scales as 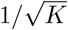 (Fig 1E). The scaling is different for high variance where *f* scales closer to 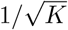 (Fig 1D) and *μ* is relatively independent of *K* (Fig 1E). Thus, the high-variance case, which we call sparse balance, results in networks in which activity is sparse but individual units have appreciable responses.

A well-known distinctive feature of standard balanced networks (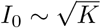 and *J*-variance ~ 1*/K*) is that the average firing rate 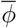 is a linear function of the bias input *I*_0_ despite the presence of a nonlinear response function in the model. This feature extends to the low bias model (*I*_0_ ~ 1) in the case of low variance but, for high *J* -variance 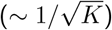, the average response 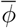 has a nonlinear dependence on *I*_0_ (Fig 1F). In both the low- and high-variance cases, *f* is insensitive to *I*_0_ (Fig 1G), meaning that the mean firing rate of the active units *μ* is also linear for low variance and nonlinear for high variance (Fig 1H). Thus, the restriction to linear responses for uniform input does not apply to the sparse balance networks.

We also examined the distribution of *x* values in these networks (Fig 2). In the low-variance case, these distributions are Gaussian, and both their mean and variance decrease with *K* (Fig 2A). The result of these two effects is that the fraction of the *x* distribution above threshold (*x* = 0) remains fairly constant as a function of *K*, corresponding to the roughly constant fraction of active units (Fig 1D), but the range of the distribution above threshold drops with *K*, matching the drop in activity (Fig 1E). For high variance (Fig 2B), the distribution is non-Gaussian, the fraction above threshold drops with *K*, and the range remains constant, again corresponding to the dependence of the average firing-rate response on *K* (Fig 1D & E). The mean of the *x* distribution for the sparse balance network is insensitive to *K* and lies below threshold. The mean of the distribution for low variance is also negative, but it moves toward zero as *K* increases.

**Fig 2.**
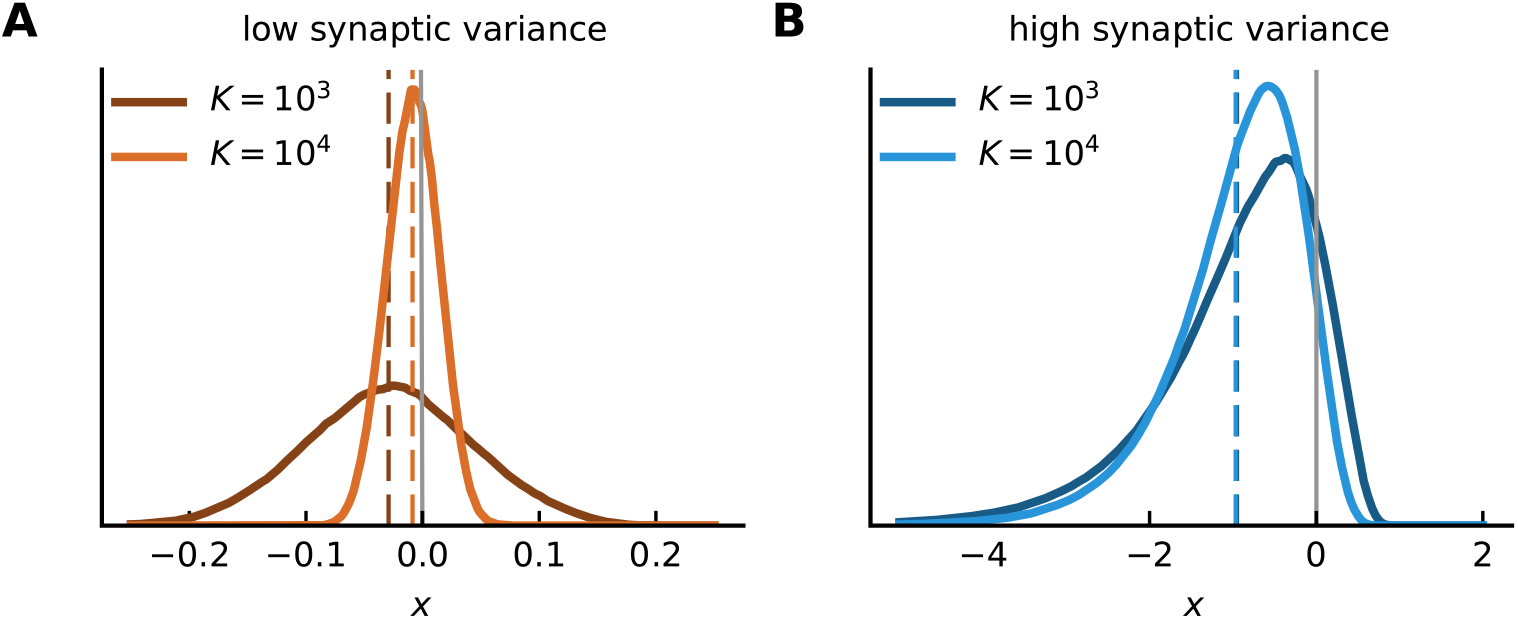
Sparse balance yields non-Gaussian dynamics and a subthreshold mean. Distribution of currents *x* (over time and units) for gamma-distributed synapses. Dashed lines denote the mean of each distribution, i.e., 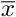. Area above threshold (set to zero; solid line) corresponds to the fraction of active units *f*. **A)** With low synaptic variance (*ν* = 1), the distribution of *x* is a Gaussian centered around a mean that tends to zero for larger *K*. **B)** Same as in A except for high synaptic variance (*ν* = 1/2). Note the larger range of the horizontal axis compared to B. The distribution is no longer Gaussian. 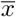 is relatively insensitive to *K* and lies below threshold. (Model parameters are *g* = *J*_0_ = 2, *I*_0_ = 1, *J_ij_* ~ gamma, *ϕ* = [tanh]_+_)

The results for networks with small input biases and large synaptic-weight variances, shown in Fig 1 for a rectified hyperbolic tangent nonlinearity, extend to other nonlinear response functions as well (Fig 3A-B). The response in these networks is distributed across almost all of the units, but at any given time only a sparse distinct subset of units is active (Fig 3C). This active population constantly changes, and firing rates appear chaotic. The fraction of time that units are active is skewed toward small values (Fig 3D), indicating that the majority of units respond infrequently. For all choices of *ϕ*, the distribution of *x* is non-Gaussian with only a small fraction of units above threshold (Fig 3E), consistent with the sparsity of the firing. Finally, the dynamics in these networks can be characterized by the population-averaged autocorrelation function of the currents (Fig 3F), which we consider in more detail in a later section.

**Fig 3.**
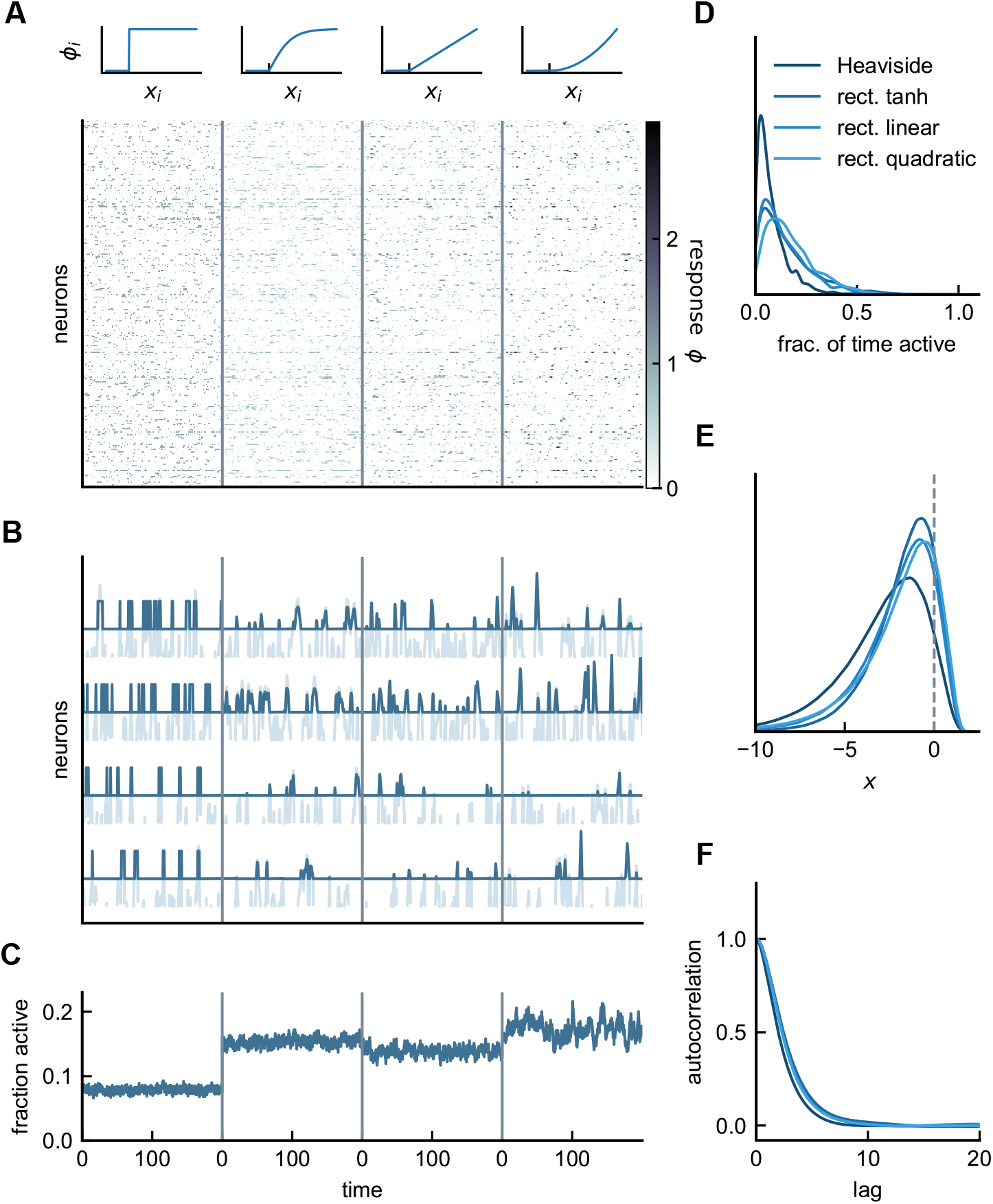
Asynchronous irregular activity in the sparse balance model. **A)** Responses of network neurons in time for four different nonlinear response functions: Heaviside step function, rectified tanh, rectified linear, and rectified quadratic. **B)** Rates *ϕ*(*x*) (dark) superimposed on the currents *x* (light) for four example units. Cells respond robustly and infrequently across choices of the response functions. **C)** Fractions of active neurons, or the inverse sparsity. **D)** Normalized distributions for the fraction of ON-time, defined as the fraction of (simulation) time a unit spends above threshold. For better visualization, histograms are smoothened using kernel density estimation. **E)** Normalized distributions of *x*, showing non-Gaussian dynamics. **F)** Population-averaged autocorrelation functions of *x*. At this fixed value of in-degree (*K* = 1000), all response functions produce qualitatively similar results. (Model parameters: *g* = *J*_0_ = *I*_0_ = 2, *J_ij_* ~ gamma).

In summary, these simulations illustrate an alternative regime for E-I networks in which the activity of individual units remains robust, despite the absence of order 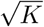 feedforward bias inputs. Furthermore, in these networks, mean firing rate exhibits a nonlinear dependence on bias input. We now analyze in detail the features illustrated in these simulations.

### Analysis of sparse balance networks

How does high-variance connectivity support sparse but robust activity with low bias input, and what is the nature of this activity? Addressing these questions is simplified by considering a Heaviside response function (Eq (3) with *λ* = 0; we comment on extensions to other nonlinearities in the Materials & Methods). For a Heaviside nonlinearity, the firing rate of an active unit is always one, so *μ* = 1 and the average response is equal to the sparsity, 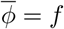. We consider a general *J* -variance scaling, 1*/K^ν^*, so that we can compare results to the low-variance *ν* = 1 case, but we are primarily interested in the high-variance case *ν* = 1/2.

We begin the analysis by defining the recurrent synaptic input as

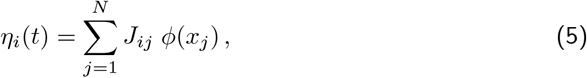

so that Eq (1) can be written as

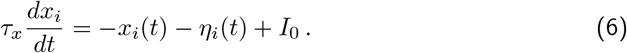

We consider non-zero weights drawn from a gamma distribution, gamma(*κ, θ*), where *κ* and *θ* are the ‘shape’ and ‘scale’ parameters of the gamma distribution in terms of which its mean is *κθ*, and its variance is *κθ*^2^. To achieve a mean 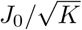 and variance *g*^2^*/K^ν^* we set

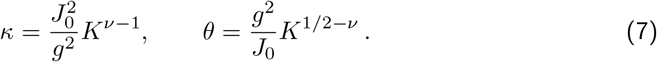

For a Heaviside nonlinearity, the sum in Eq (5) is only over active units with *ϕ* = 1, and the probability of a unit being active is equal to the sparsity *f*. This means that of the *K* non-zero elements of *J* for each unit, *f K* will be active. As a result, *η* is given by the sum of *f K* random variables drawn independently from the distribution gamma(*κ, θ*). The sum of random variables that are gamma-distributed with the same scale parameter is itself gamma-distributed with that scale parameter and a shape parameter equal to the sum of the shape parameters of the variables being summed [18]. Thus,

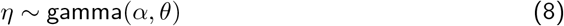

with *α* = *f Kκ*, which has mean

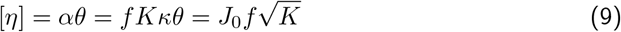

and variance

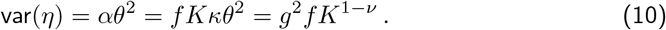

To maintain a finite mean input as *K* grows, Eq (9) implies that we must have 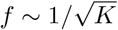,which implies, from Eq (10), that the fluctuations in the synaptic input scale as *K*^1/2*−ν*^. Thus, the only solution with finite fluctuations as *K* grows is the high-variance case, ν = 1/2. For ν = 1/2 and with 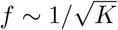, Eqs (9) and (10) show that the distribution of synaptic inputs is independent of *K* with 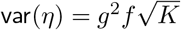. This feature may appear surprising given that the sparseness of network activity is proportional to 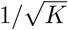. We resolve this paradox in a later section.

A naive application of the central limit theorem would suggest that for sufficiently large *K*, the synaptic input would be normally distributed. Independent of *θ*, the larger the shape parameter, the closer a gamma distribution approximates a Gaussian (in particular, the approximation is good for shape parameters ~20 or larger). The shape parameter for the distribution of *η*, from Eq (8), is 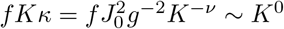. Thus, unless *J*_0_ is large or *g* is small, even in the limit of large *K*, the *η* distribution remains non-Gaussian (Fig S4).

If we average Eq (6) over both units and time and use 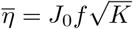, we obtain 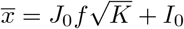 or, equivalently,

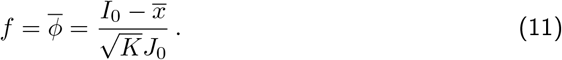

In the standard balanced model and in the low-variance case considered above, 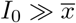, so the mean response is linear in *I*_0_. This is no longer true for high synaptic variance (*ν* = 1/2) for which *I*_0_ and 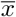 are both of order 1. The nonlinear mean response seen in Fig 1F arises because the dependence of 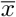 on *I*_0_ is nonlinear. Depending on the choice of *ϕ*, the sparse balance model exhibits sublinear or subralinear mean population response (Fig S5).

These analyses show that when feedforward bias input is of order one, large synaptic variance is required to generate robust fluctuations, with a synaptic variance of order 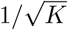 producing order-one fluctuations.

### Sparse activity arises from network dynamics

We noted in the previous section that the distribution of synaptic inputs is independent of *K*, and yet the mean network firing rate 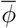 varies as 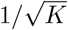. Network currents *x* are generated through Eq (6), which involves low-pass filtering of the synaptic input. This suggests that the response sparseness is related not to the distribution of synaptic inputs but rather to their dynamics.

To explore these dynamics, we consider the population-averaged autocorrelation function of *η*,

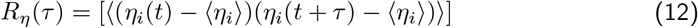

which captures the extent to which *η* at time *t* + *τ* is affected by *η* at time *t*. *R_η_* is a decaying function of the lag *τ* (Fig 4A), and its decay defines a correlation time-scale denoted by *τ_η_*. One way to define this correlation time-constant is by considering the normalized area underneath the autocorrelation function,

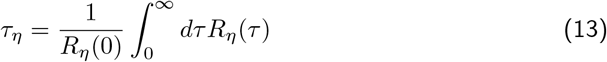

**Fig 4.**
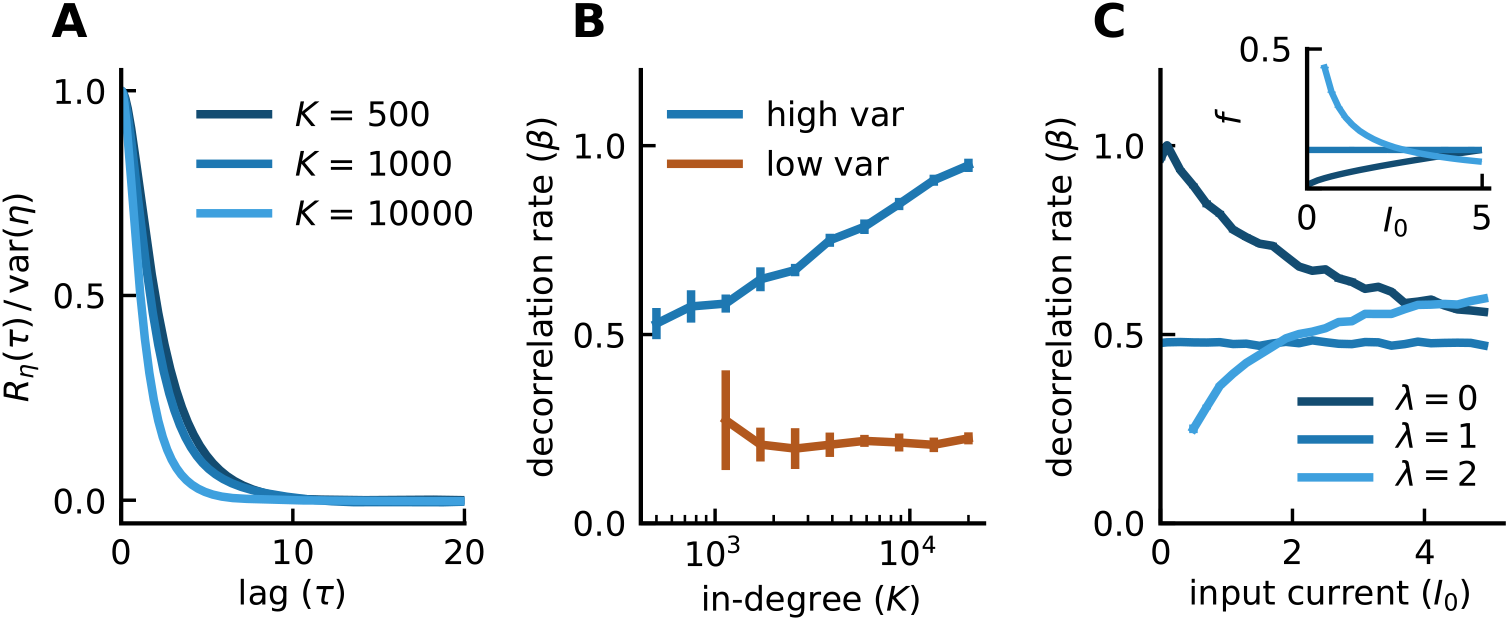
Time-scale of fluctuations adjusts to maintain sparse activity. **A)** Population-averaged autocorrelation function of the synaptic input normalized by its zero-lag value. Note the faster decay of the autocorrelation for increasing *K*. **B)** The decorrelation rate *β* is constant in the low-variance network but increases logarithmically with *K* in the sparse balance model, resembling the (inverted) trends of sparsity (Fig 1D). **C)** *β* also exhibits a nonlinear dependence on *I*_0_ similar, but opposite to, that of the sparsity (inset). Error bars indicate the 95% confidence interval around the mean, averaged over 10 random realizations of the connectivity. (Model parameters: *g* = *J*_0_ = 2, *I*_0_ = 1, *K* = 1000, *J_ij_* ~ gamma, 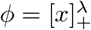 for *λ* sweeps in **C**, [tanh]_+_ otherwise.)

We characterize the dynamics of *η* using the dimensionless constant *β* = *τ_x_/τ_η_*, and find that *β* increases logarithmically with *K*, an increase that does not occur in the low-variance (Fig 4B) or for conventional balanced networks. As *K* increases, the time-scale of the fluctuations in *η* becomes more rapid, although their variance remains constant. This makes it increasingly harder for *x* to keep up with the fluctuations, due to the low-pass filtering in Eq (6). As a result, the fraction of *x* above threshold decreases and the overall activity decreases with *K*. Thus, interestingly, it is the dynamics of the recurrent synaptic inputs, not their size, that leads to sparse activity at large *K*.

Consistent with the argument above that links the time-scale of dynamics to the level of currents above threshold, we find that changes in *β* account for the degree of sparsity (Fig 4C). In particular, trends in *β* are opposite to those in *f*, with sparser activity (smaller *f* ) corresponding to faster time-scale (larger *β*) and vice versa. These results highlights how the time-scale of synaptic fluctuations dynamically adjust to maintain sparse activity.

### Mean-field analysis

In a previous section we noted that, for a Heaviside response nonlinearity and gamma-distributed synaptic weights, the recurrent synaptic input *η* is gamma distributed with shape parameter (for the high-variance *ν* = 1/2 case) 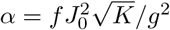 and scale parameter *θ* = *g*^2^*/J*_0_. Although *θ* is completely determined by parameters characterizing *J* (*g* and *J*_0_), *α* is not determined because it depends on the fraction of active units *f* or, equivalently, on the mean firing rate 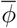. We begin our mean-field analysis by deriving a self-consistent equation for the mean response that determines *α* and, thereby, the full distribution of recurrent synaptic inputs. For this purpose, we introduce the variable *m*, which is the mean-field approximation for 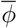. The above equations imply that *α* is given in terms of *m* by

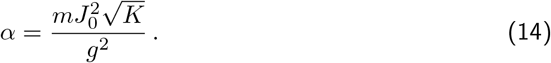

Our goal is therefore to compute *m* as a function of *α* so that the above equation becomes a closed self-consistent condition for determining *α*.

Conventionally, in a dynamic mean-field approach, the full autocorrelation of *η* is computed self-consistently [15, 16, 19, 20]. This computation is difficult in the high-variance case because of the non-Gaussian statistics of *η*. Instead, we consider a ‘static’ mean-field approximation motivated by the logarithmic scaling of the decorrelation rate *β* in the sparse balance model. When *β* is small, *x* roughly tracks the slow fluctuations in *η*. This tracking holds particularly well for *β* ≪ 1. On the contrary, for *β* ≫ 1, *x* cannot keep up with the fluctuations in *η* and averages them. As Fig 4B suggests, for in-degrees of interest, *K* ~ 10^3^, the network operates closer to the tracking regime, allowing us to make the approximation *x* ≈ −*η* + *I*_0_. Indeed, for *K* ~ 500−1000, the distribution of *x* resembles that of *η*, except for a rightward shift by *I*_0_ (Fig 5A). Note that this approach differs from mean-field approaches in which the limit *K* → ∞ is assumed. Here *K* is considered large but finite. In both cases, however, the limit *N* → ∞ is assumed.

When *x* tracks *η*, we can use the distribution of *η* values, *η* ~ gamma(*α, θ*), to perform averages over *x*. For example, for the mean-field calculation of [〈*ϕ*(*x*)〉], we can make the substitution *ϕ*(*x*) → *ϕ*(*I*_0_ − *η*) and write

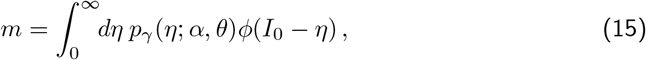

where *p_γ_* is the probability density function of the gamma distribution with shape and scale parameters *α* and *θ*. Eqs (14) and (15) together form a closed self-consistent condition that determines *α*, with results that are in decent agreement with numerical simulations (Fig 5B). In particular, the nonlinear relationship between *α* and *I*_0_ is captured by the theory. Larger values of *K* exhibit slightly larger deviations from the theory, which hints at the violation of the static assumption. However, even with *K* ~ 10^3^, which is in the range of interest, the theoretical *α* is close to the empirical results (Fig 5B).

**Fig 5.**
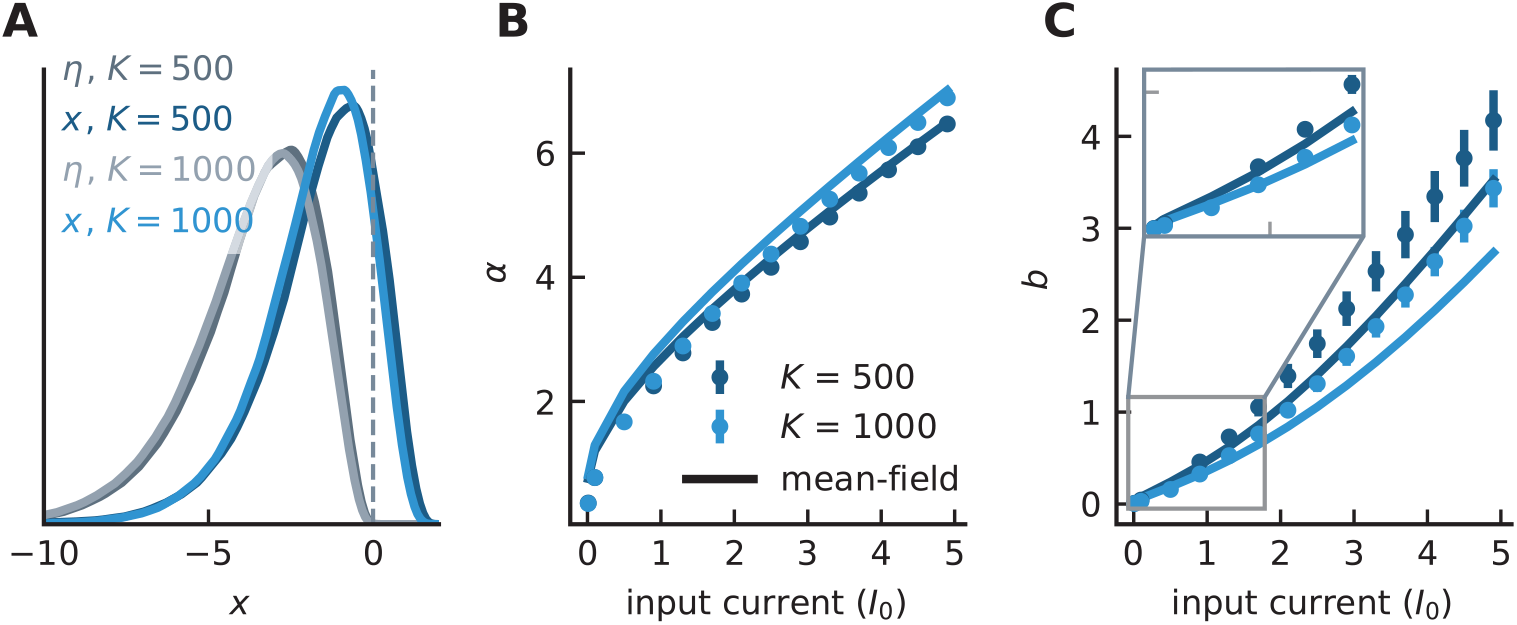
Mean-field approximation captures the mean and quenched variance of the synaptic input. **A)** Distribution of the recurrent synaptic input *η* (gray) and current *x* (blue) for two *K* values. **C-D)** *α* and *b* as functions of input current *I*_0_. The nonlinear features of *α* and *b* are captured by the theory. The approximation for *b* works better with small currents (zoomed-in gray box). The trend (larger *K* produces smaller quenched variance and larger *α*) is also accounted for by the theory. Error bars indicate the 95% confidence interval around the mean, averaged over 10 random realizations of the connectivity. (Model parameters: *I*_0_ = 2 in **A**, *g* = *J*_0_ = 1, *J_ij_* ~ gamma, *ϕ* = Heaviside)

Although we have computed *α* and thereby determined the distribution of *η* values, this does not completely characterize the nature of the fluctuations in the recurrent synaptic input. The total variance of the recurrent synaptic input can be divided into temporal and quenched parts by writing

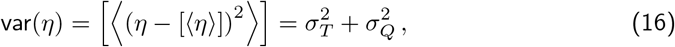

where

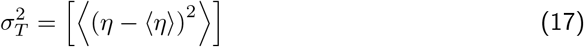

and

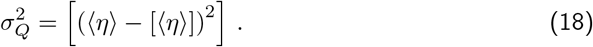

The quenched variance arises because the different units of the network fluctuate around different time-averaged values. To analyze this quenched variance, we need to specify how the total variance is divided into temporal and quenched components. For this purpose, we decompose *η* as

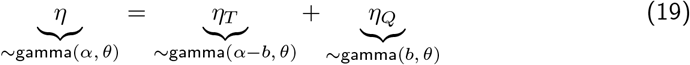

where *η_T_* is a time-dependent variable with no quenched variance, and *η_Q_* is a static variable that embodies the influence of quenched disorder. The shape parameters of the temporal and quenched components must add up to *α*, and the variance of the quenched component determines *b* through

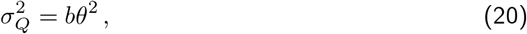

where *b* is the scaled quenched variance. This decomposition assumes that the time-averaged synaptic input in the full model is gamma-distributed (Fig S6).

Our mean-field analysis of the quenched variance, *b*, is based on computing a mean-field approximation, *s*, of [〈*ϕ*〉^2^]. Using the decomposition into temporal and quenched components of the gamma distribution, we can write the mean-field approximation of *s* as

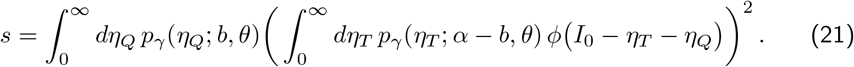

This expression depends on the parameter *α* computed above and, in addition, on *b*, so we need a self-consistency condition to determine the value of this second parameter. Using standard mean-field approximations, we can write

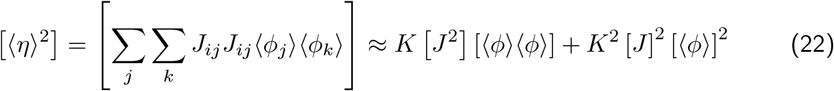

Inserting now 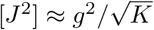 and 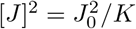 gives

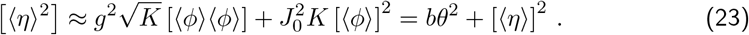

Finally, using 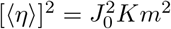 and [〈*ϕ*〉] = *m*, this yields

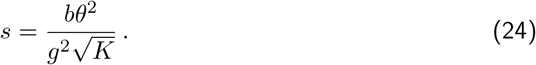

Eqs (21) and (24) form a closed system that can be used to determine *b* (Materials & Methods). Fig 5C depicts this self-consistent solution. Notably, the theory captures the nonlinear relationship between *b* and *I*_0_ as well as the trend with *K*. Since the mean-field solution to *α* is used in computing *b*, any error in the estimate for *α* is carried over to the solution for *b*. The primary source of error is the violation of the assumption that the distribution of time-averages are gamma-distributed, which is used in the decomposition of *η* in Eq (19). The deviation is pronounced for larger bias inputs but for *I*_0_ ~ 1, which is closer to the range of input currents of interest, the theory is in decent agreement.

These results suggest that, despite non-Gaussian statistics, the sparse balance model is amenable to a mean-field treatment that is in similar spirit to what has been applied previously to recurrent networks.

### Sparse balance in an E-I network

Finally, we illustrate that all of the features we have discussed for a purely inhibitory network are present in mixed excitatory-inhibitory networks for two choices (gamma and lognormal) of the connectivity distribution (Fig 6). These networks exhibit asynchronous irregular activity with chaotic responses of individual units (Fig 6A-B) and constant population activity (Fig 6C). Responses are sparse across the population: roughly 10% of excitatory and 20-30% of inhibitory units are active at any given time (Fig 6C). Individual units show sporadic response for both E and I cells in time (Fig 6B) with spatiotemporal variability that is purely internally generated.

**Fig 6.**
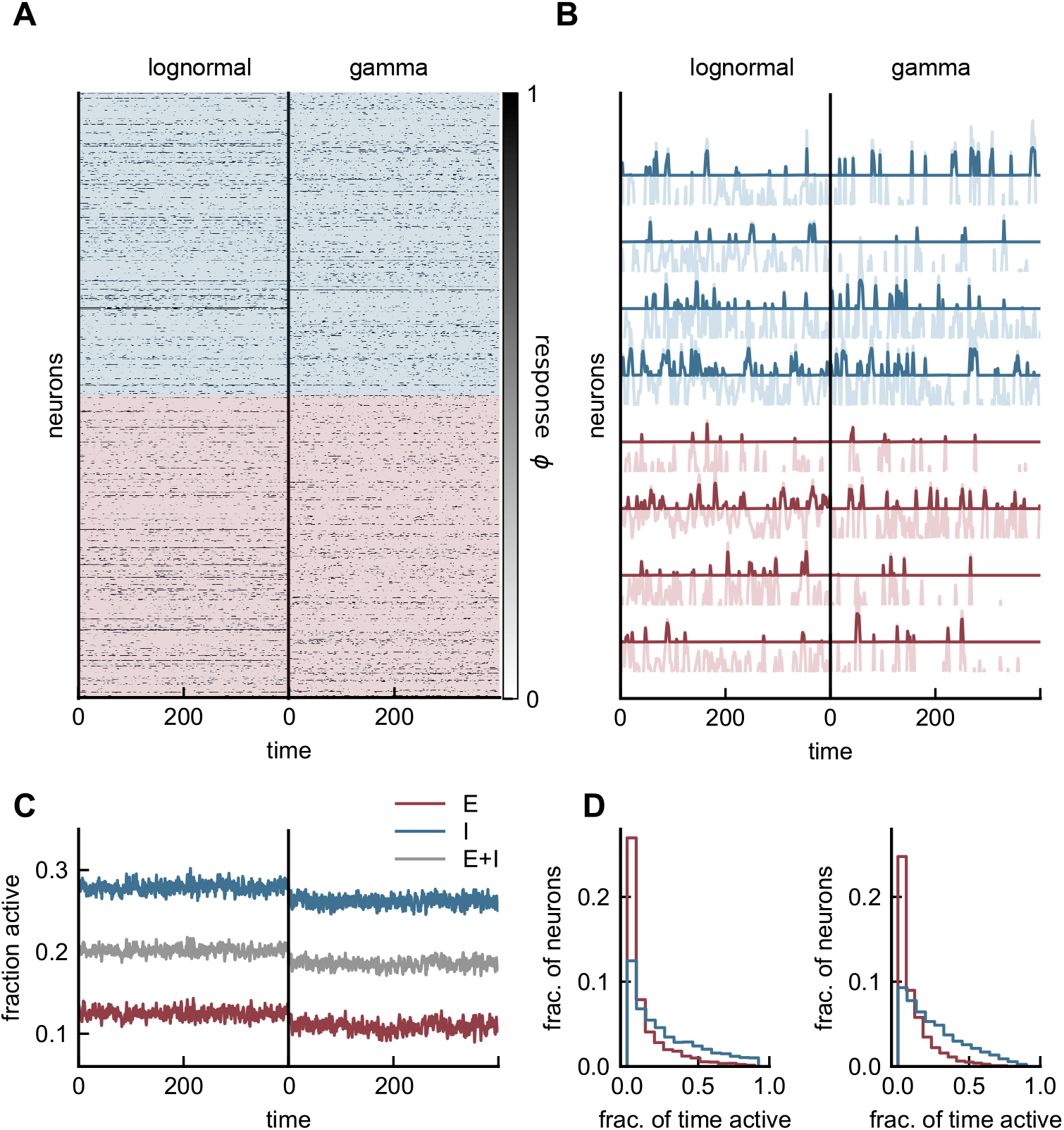
Asynchronous irregular activity in an E-I network with small input current. **A)** Responses of 500 excitatory (red) and 500 inhibitory (blue) units in two networks, one with lognormal (left) and the other with gamma (right) weight distributions. Responses are sparse and distributed across the population. **B)** Rates (dark) superimposed on the currents (light) for four example cells from each population. Response is infrequent as fluctuations occasionally push the current above threshold. **C)** Fraction of active units for individual populations (red and blue) and across the entire network (gray). The inhibitory population is more active than its excitatory counterpart. **D)** Fraction of (simulation) time units spend above threshold for each population and connectivity distribution. This distribution is wide and skewed. Both choices of the connectivity distribution produce qualitatively similar results. (Model parameters: *g* = 1, *J_EE_* = *J_IE_* = 1, *J_EI_* = 2, *J_II_* = 1.2, *I_E_* = 2, *I_I_* = 1, *N_E_* = *N_I_* = 3000, *K* = 600, *ϕ* = [tanh]_+_).

Responses are robust and shared across the entire population as opposed to a fixed subset of units. We characterize this feature by considering the distribution of ON-time fraction, i.e., the fraction of time individual cells spend above threshold (Fig 6D). This quantity shows a wide and skewed distribution across both E and I populations. The majority of units spend very little time above the threshold, with only a few (5% of E, 20% of I cells with lognormal; 2% of E, 15% of I cells with gamma) spending more than half the time above threshold, and none responding at all times. We note that gamma and lognormal synaptic distributions produce similar activity patterns across the population.

## Discussion

We have uncovered a novel regime of E-I networks that exhibits asynchronous irregular activity without the need for unrealistically large external input currents. We have done so by taking advantage of widely distributed synapses that generate fluctuations that would otherwise be minuscule in the absence of large feedforward currents. We highlighted a number of properties including sparse activity, non-Gaussian dynamics and a nonlinear population response. We also revealed the mechanism by which the time-scale of the dynamics generates sparse network activity. Using mean-field theory, we computed the statistical features of the recurrent input. This model demonstrates the important role of synaptic variance in the dynamics of recurrent networks.

Robust network responses with small input currents are especially interesting in light of the fact that experiments suggest the feedforward component of the input in cortical circuits is comparable in magnitude to the total synaptic input (see [9] for a review). For example, in olfactory cortex, the feedforward excitation from the olfactory bulb accounts for only a quarter of the net excitation into a pyramidal cell [10]; in the visual cortex, thalamic input accounts for roughly 30-40% of the net excitation to a cell [11, 12]. Similar results have been reported in the auditory cortex [21]. Our model provides a novel theoretical insight into these observations and highlights the importance of the scaling of the input current on network dynamics.

To provide a more quantitative link to these experimental findings, it is appropriate to define 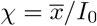, referred to as the ‘balanced index’ [9]. The ratio *χ* captures the relative contribution of the feedforward input *I*_0_ to the mean of the total current 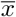. The aforementioned experiments suggest a *χ* of order 1. In both the standard balanced and the low-variance networks result in 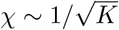. In the sparse balance model, widely distributed synapses together with small input currents yield a *χ* of order 1 in agreement with experimental findings in cortex.

Despite the absence of cancellation of large excitatory and inhibitory currents in the sparse balance model, the mean of the net synaptic input, 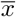, lies well below threshold. One consequence of this, as mentioned above, is that the net input and the feedforward input have comparable contributions to the mean response. This results in a nonlinear population response to uniform input that is absent in the standard balanced regime where the strength of the feedforward input overwhelms the influence of the net input. Nonlinear mean responses are known to be necessary for a variety of cortical computations such as response normalization and surround suppression in visual cortex [9, 17, 22–24] and concentration invariance in olfactory cortex [25–27]. In the sparse balance model, the shape of this nonlinear population response depends on the choice of neuronal response function.

Small input currents impose a constraint on the mean response. To ensure this constraint is carried over to the sparsity, but not the mean response of the active neurons, we considered widely distributed synapses through an unconventional scaling of synaptic variance. Models that address the role of heavy-tailed connectivity distributions are timely because it has been shown that the distribution of synaptic efficacies in cortex are compatible with a lognormal fit [1, 28, 29]. Experiments and modeling studies have also suggested that strong synapses in the tail of such distributions, although less frequent, can have a strong influence on postsynaptic firing and network dynamics [30–33].

Our choice of the gamma distribution, as opposed to the lognormal, was motivated by its analytical tractability. The scaling of variance we consider results in an effectively sparse connectivity distribution where the majority of synapses are weak and thus neuronal activity is heavily influenced by the minority of strong synapses. Similar to this idea, recent modeling work has demonstrated that networks with power-law synaptic weights exhibit self-sustained activity [33]. In these ‘Cauchy networks’, the variance of the connectivity is infinite for finite *K*, and network behavior is dominated by large tails in the weight distribution. In another modeling study, a lognormal distribution of synaptic weights in a network of spiking neurons gave rise to self-sustained asynchronous firing in the absence of any bias input current [31]. Together with these results, our model highlights the degree to which heterogeneity in connectivity can compensate for the absence of large input currents and help sustain rich network dynamics.

The emergence of sparse activity in our model is interesting given that only a small fraction of cortical cells, particularly in the superficial layers, are active in response to many stimulus or spontaneously (see [34] for a review). The level of sparsity in the model is related to the degree of network connectivity as opposed to single-neuron properties, such as the threshold. Standard balanced models can also exhibit a high degree of population sparseness, as in the spiking models of [35]. In networks with intrinsic chaotic activity, whether and how the degree of sparsity can be used to perform computations that require high-dimensional representation [36, 37] of the stimulus remains to be investigated.

In our model, we revealed that the time-scale of the synaptic input, not its distribution, adjusts to maintain the degree of sparsity in the network. When the constraint on the mean response demands a small level of current above threshold, fluctuations at the synaptic input speed up; since the dynamic equation of each cell is a low-pass filter of its synaptic input, the output distribution narrows ever so slightly and the tail above threshold retreats in a manner that satisfies the constraint on the mean rate. Changes in sparsity are made possible by changes in the autocorrelation time-scale, and this phenomenon appears to apply generally. This feature highlights the flexibility of recurrent dynamics in adjusting not only the mean and variance of the distribution of its firing rates, but also their correlation time-scales. The role of this flexibility in building functional networks of rate neurons is a potential subject of future work.

Mean-field theory is an important tool for understanding network behavior. We examined a mean-field theory based on the approximation that synaptic fluctuations are instantaneously tracked by neuronal responses. This adequately predicted the network behavior and was extended to include both time-dependent and quenched fluctuations. This theory assumes a finite, but large in-degree *K*. Biological values of *K* are roughly of order 10^3^, the range we considered. Specifically, pyramidal cells receive ~7000 excitatory synapses [1], but when we account for the average number of synapses per connection, ~4 [38, 39] and the number of non-intracortical synapses, the in-degree comes out to be ~10^3^.

One prominent theory that addresses the issue of bias input is the stabilized supralinear network (SSN) [9, 24, 40]. Important features of SNNs include supralinear neuronal response function, small bias current, and weak synaptic coupling. These models have been extensively and successfully used to describe steady-state responses in sensory cortex. Our model differs from the SSN in that it generates chaotic activity, has strong synaptic coupling [41] and has widely distributed synaptic weights. With a supralinear response function (*λ* > 1), the large synaptic variance endows our model with a higher degree of chaos than other strongly-coupled networks; without this large degree of heterogenity in the weights, responses are susceptible to resting at fixed points [15]. This degree of chaos may aid in the learning of functional trajectories [42–46].

We examined a model with random connectivity, but it would be interesting to investigate stimulus selectivity in sparse balance networks with structured connections. The large degree of variability in the synapses could route stimulus information along particular paths across network neurons. Structured connectivity is of particular interest given compelling evidence that the recurrent contribution of the synaptic input, not just the feedforward component, exhibits selectivity [10–12]. We believe that the variance of connectivity, in addition to its mean structure, is important to consider for addressing the way feedforward and recurrent components shape selective responses.

## Materials & methods

### Numerical simulations

Numerical simulations were performed using Euler integration with time-steps less than 0.05, *τ_x_* = 1, and simulation time *T* = 1000. Other network parameters are included in the figure captions. The code (written in Julia v1.3.0) is available with the online version of the manuscript.

### Non-Heaviside nonlinearities

Our analysis, leading to the high variance scaling (*ν* = 1/2) is based on networks with a Heaviside (*λ* = 0) response function, which simplifies the calculations. Generally, the variance of the recurrent synaptic input is 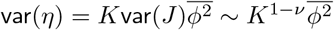. With a Heaviside nonlinearity, 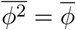 and since 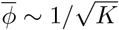, we conclude that *ν* = 1/2 is the only solution with finite fluctuations as *K* grows. Non-binary responses do not guarantee the equivalence of 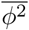 and 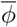, so we introduce 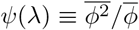. In the case of a Heaviside (*λ* = 0), ψ = 1. For *λ* > 0, the distribution of *x* dictates this ratio. We find numerically that, while not exactly one, *ψ* is slowly varying in *K* (Fig S3).

The variance scaling result obtained with a Heaviside nonlinearity extends to rectified tanh (and similarly rectified linear) in that the 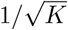 scaling of 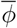 is predominantly inherited from *f*, and not *μ*. For *λ* ≥ 2, this is not necessarily the case and *μ* can exhibit a non-negligible scaling with *K*. In spite of this, with a fixed value of *K* in the range of interest, ~10^3^, the responses in the high-variance case, as opposed to its low-variance counterpart, are much more appreciable and irregular, and the low-variance model is prone to fixed point states for *λ* > 1.

### Existence of the mean-field solution

From Eq 15,

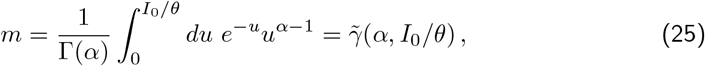

where 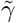 is the regularized lower incomplete gamma function. From Eq 14,

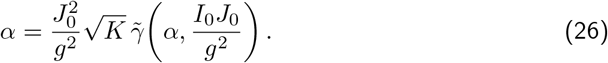

Solving this equation for *α* produces the result shown in Fig 5B.

Combining Eqs (21) and (24), we obtain

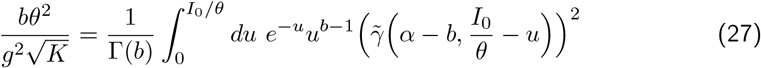

Here, we comment on the existence of a solution to Eq (27). On one hand, the maximal possible value of *b* is *b* = *α*. At this extremum value, it follows from Eq (14) that the left-hand-side (LHS) of Eq (27) is equal to *m*. In this limit, the right-hand-side (RHS) of Eq (27) is also equal to *m*. This is because 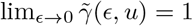. In other words,

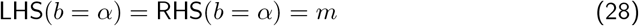

This, however, is not a viable solution because *b* = *α* describes a fixed-point state with no temporal variability, rather than describing a chaotic state.

On the other hand, the minimal value of *b* is *b* = 0, and a similar manipulation of Eq (27) shows that RHS(*b* = 0) = *m*^2^

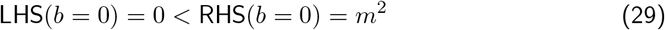

Therefore, a solution is guaranteed as long as the slope of the RHS at *b* = *α*, *∂*(RHS)*/∂b*, is larger than that of the LHS. These slopes depend on *K*, through the explicit appearance of *K* in the LHS of Eq (27) and through the dependence of the mean-field value of *α* on *K*. Numerically, we find that a solution exists for the *K* values of interest. Solving Eq (27) for *b* produces the result shown in Fig 5C.

## Acknowledgments

We thank Ken Miller, Gregory Handy, Alessandro Ingrosso, and Rainer Engelken for helpful discussions. This research was supported by the NSF NeuroNex Award DBI-1707398 and The Gatsby Charitable Foundation.

---

**Fig S1.**
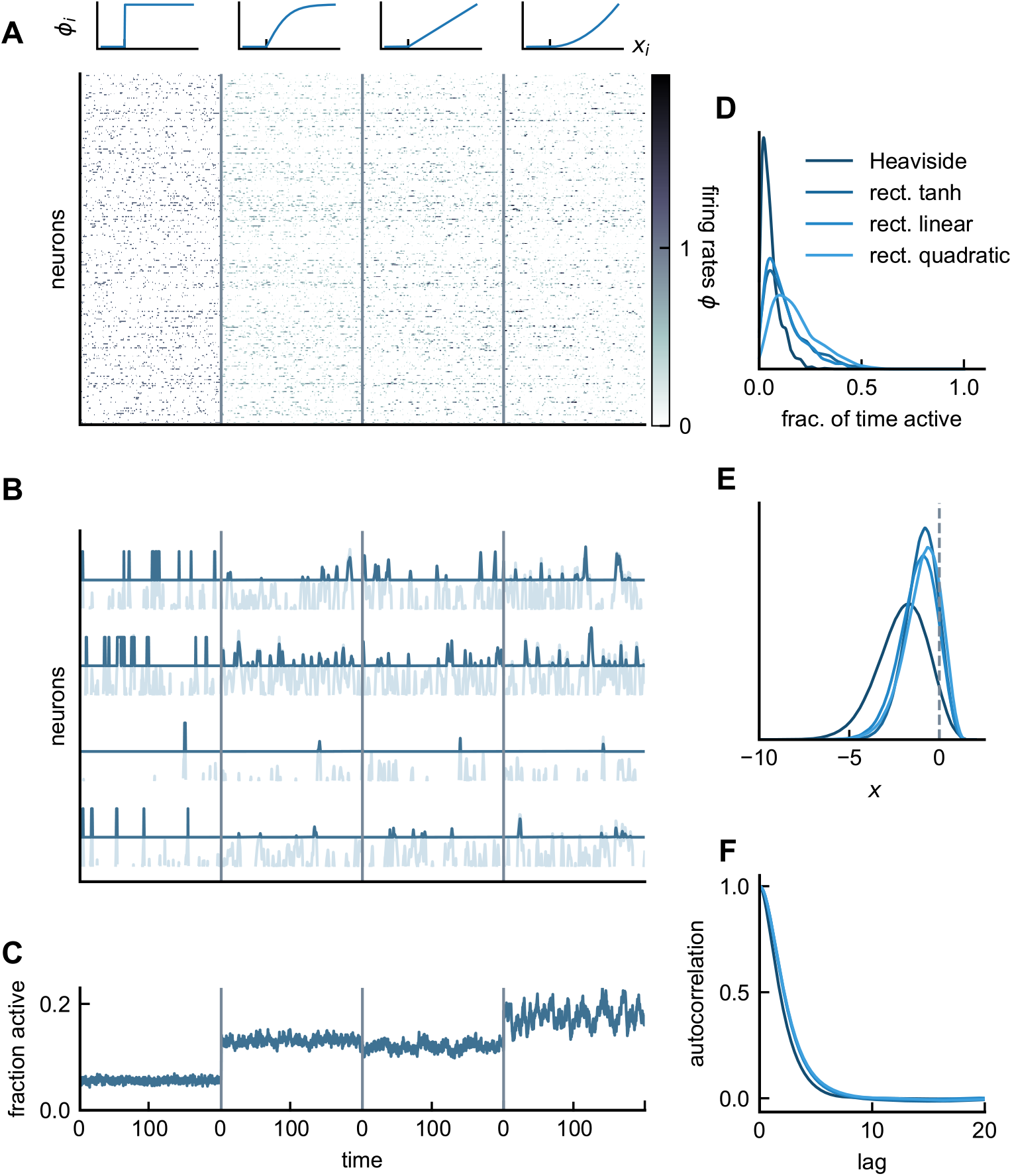
Asynchronous irregular activity in the sparse balance model with binary weights. Same as Fig 3, except with a Bernoulli connectivity distribution of mean 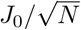. (Model parameters: *J*_0_ = 2, 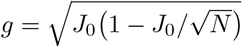, *I*_0_ = 1.5, *N* = 1000).

**Fig S2.**
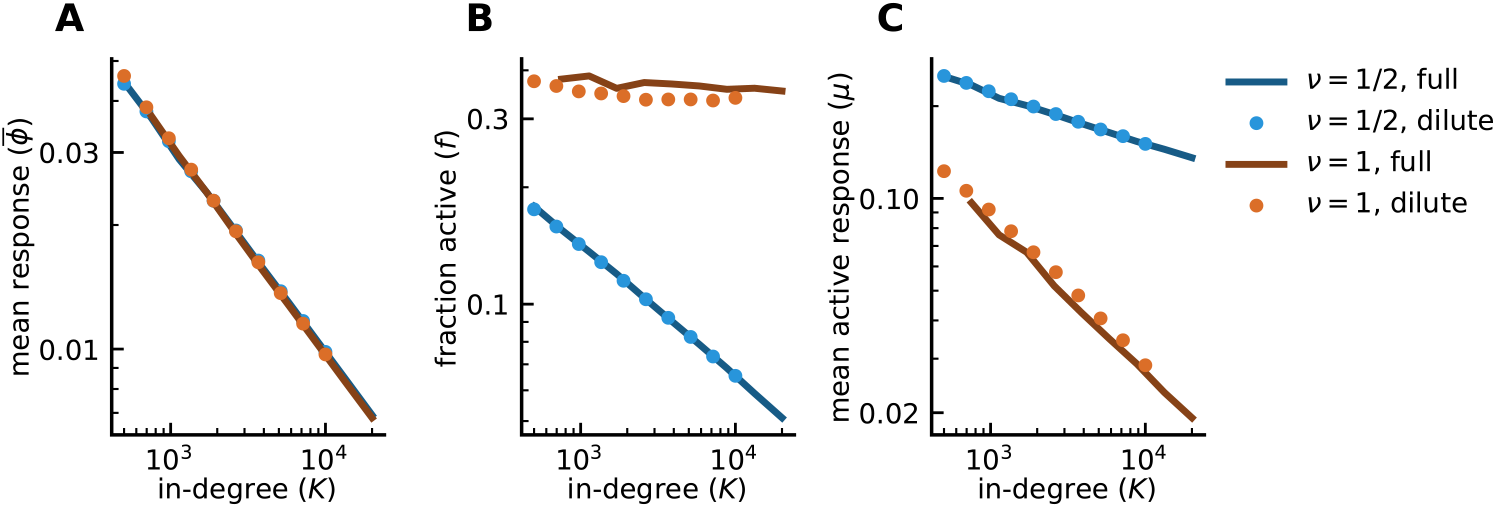
Equivalence between dilute and full connectivity. **A-C)** Same as Fig 1C-E, but with the addition of results from a dilutely-connected network (dots). With dilute connectivity, the source of variability in the connections are twofold: each neuron receives inputs from, on average, *K* other neurons out of the total *N* network neurons; additionally, each existing connection is drawn from a distribution of mean 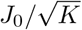 and variance 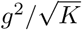. In the fully-connected case, each neuron receives input from all other neurons with connections drawn from a distribution of mean 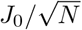 and variance 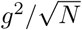. Note that in the main text, we considered fully-connected networks and denoted the mean and variance by 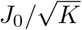 and 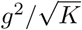 with *K* = *N*. (Model parameters: *g* = *J*_0_ = 2, *I*_0_ = 1, *J_ij_* ~ gamma*, ϕ* = [tanh]_+_; *N* = *K* in full (solid), *N* = 20000 in dilute (dotted)).

**Fig S3.**
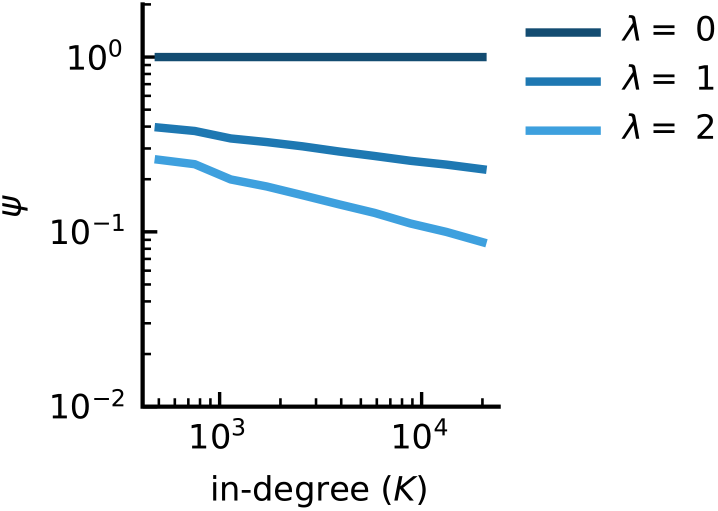
Slow variations in *ψ* for various nonlinear response functions. *ψ*(*λ*), defined as 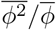, exhibits sub-power law scalings with *K*. (Model parameters: *g* = *J*_0_ = 2, *I*_0_ = 1, *J_ij_* ~ gamma, 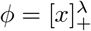)

**Fig S4.**
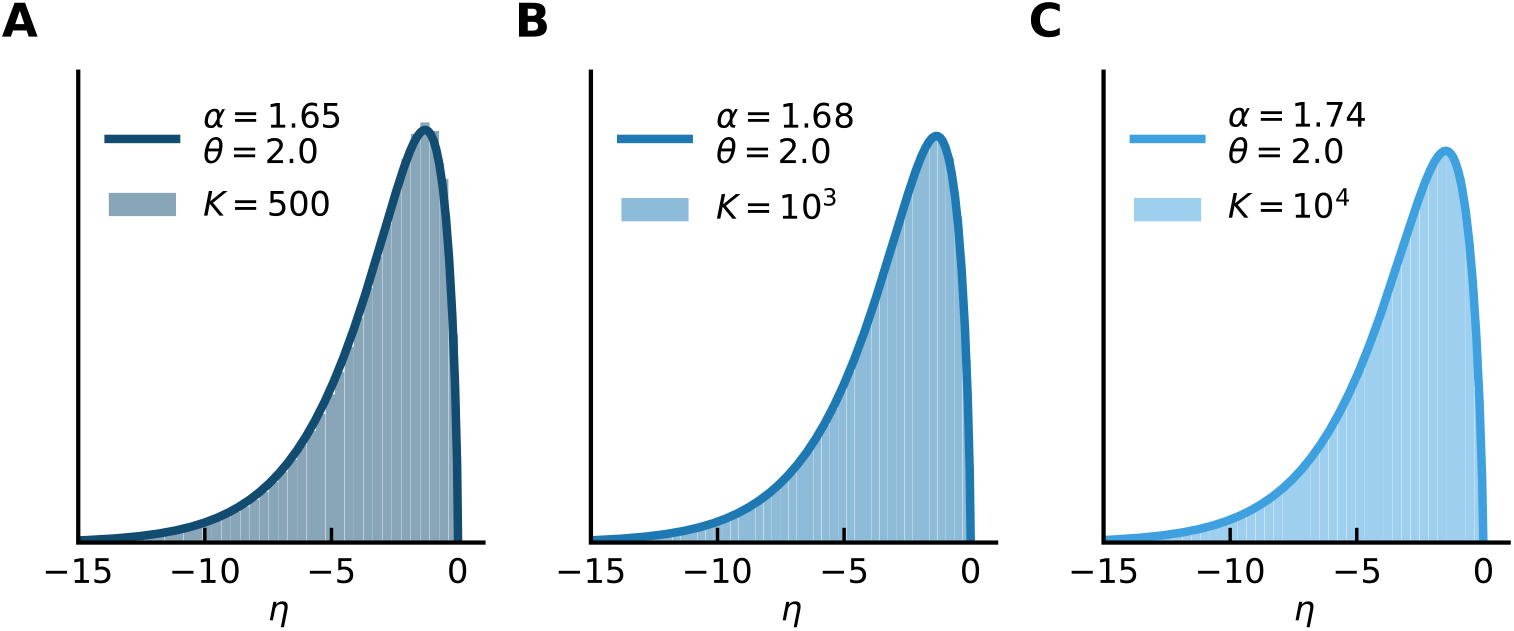
Recurrent synaptic input is described by a gamma distribution. **A-C)** The distribution of the synaptic input *η* (across population and time) with a Heaviside response nonlinearity for three different values of *K* (shaded histograms). The solid line is the gamma distribution in Eq (8) with scale parameter *θ* = *g*^2^*/J*_0_ and shape parameter 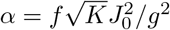, where *f* is the measured sparsity in the simulations, and *J*_0_, *g*, *K* are network parameters. The distribution accurately describes the histograms. Also note that, due to the high *J* -variance in the sparse balance model, the distributions of synaptic input hardly change as *K* increases. (Model parameters: *g* = *J*_0_ = 2, *I*_0_ = 1, *J_ij_* ~ gamma, *ϕ* = Heaviside)

**Fig S5.**
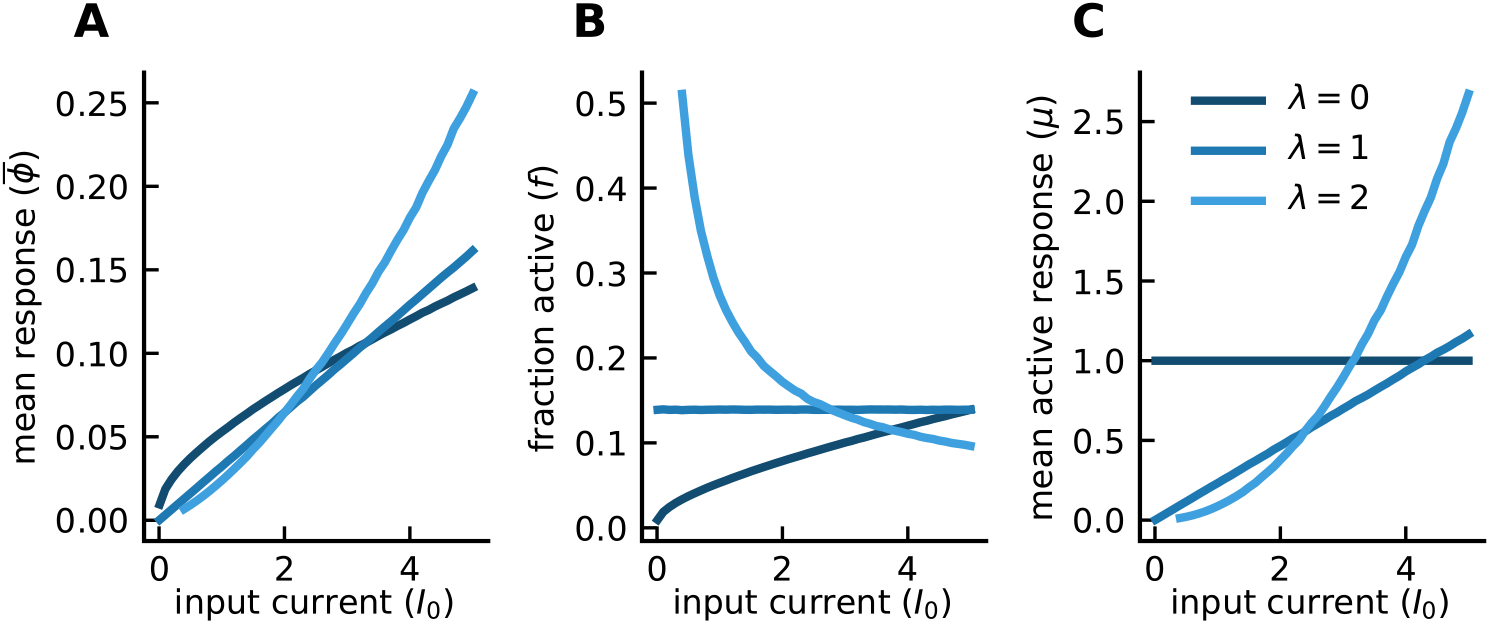
Sparse balance responds nonlinearly to input current. **A)** Mean response, 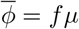, increases nonlinearly with input current. This relationship depends on the shape of the neuronal response function: *λ* < 1 and *λ* > 1 give rise to sublinear and supralinear mean responses, respectively. **B-C)** Fraction, *f*, and mean response, *μ*, of the active neurons versus input current. With rectified linear (*λ* = 1), this fraction remains constant since *x* can be rescaled by 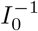 without changing the shape of the *x* distribution; this feature also makes the mean response linear. The nonlinear trend in sparsity switches at *λ* = 1. The fraction active *f* increases with *I*_0_ at an ever-decreasing rate with *λ* > 1, while the opposite is true for *λ* < 1. For *λ* > 1, with stronger feedforward excitation, threshold crossings that result in sufficiently large responses become amplified. This amplification produces a large, supralinear *μ*, which in turn comes at the cost of sparsening the population activity with *I*_0_. For the Heaviside (*λ* = 0), responses are binary, so *μ* = 1 independent of *I*_0_ and 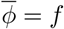. (Model parameters: *g* = *J*_0_ = 2, *J_ij_* ~ gamma, 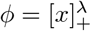, *K* = 1000)

**Fig S6.**
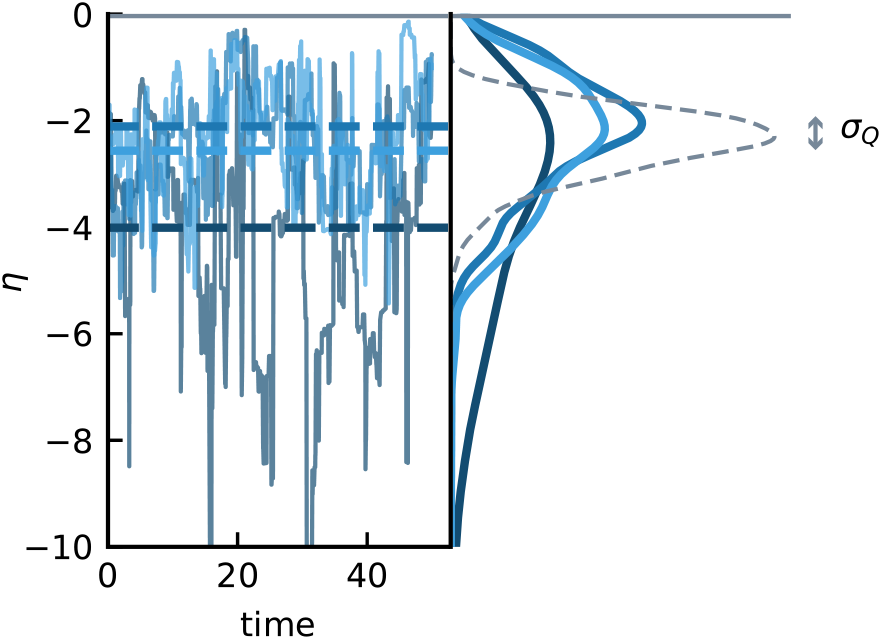
Distribution of time-averaged activities. Recurrent synaptic inputs of three example neurons with their corresponding distributions on the right (solid curves). Due to quenched disorder in the network, each neuron fluctuates around a different time-average (horizontal dashed lines). These time-averages form a distribution shown on the right (gray dashed curve) whose width is captured by 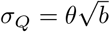 and is approximated by a gamma distribution. This approximation is the main source of error in Fig 5C. (Model parameters: *g* = *J*_0_ = *I*_0_ = 1, *K* = 1000, *J_ij_* ~ gamma, *ϕ* = Heaviside)

